# High-resolution CTCF footprinting reveals impact of chromatin state on cohesin extrusion dynamics

**DOI:** 10.1101/2023.10.20.563340

**Authors:** Corriene E. Sept, Y. Esther Tak, Christian G. Cerda-Smith, Haley M. Hutchinson, Viraat Goel, Marco Blanchette, Mital S. Bhakta, Anders S. Hansen, J. Keith Joung, Sarah Johnstone, Christine E. Eyler, Martin J. Aryee

## Abstract

DNA looping is vital for establishing many enhancer-promoter interactions. While CTCF is known to anchor many cohesin-mediated loops, the looped chromatin fiber appears to predominantly exist in a poorly characterized actively extruding state. To better characterize extruding chromatin loop structures, we used CTCF MNase HiChIP data to determine both CTCF binding at high resolution and 3D contact information. Here we present *FactorFinder*, a tool that identifies CTCF binding sites at near base-pair resolution. We leverage this substantial advance in resolution to determine that the fully extruded (CTCF-CTCF) state is rare genome-wide with locus-specific variation from ~1-10%. We further investigate the impact of chromatin state on loop extrusion dynamics, and find that active enhancers and RNA Pol II impede cohesin extrusion, facilitating an enrichment of enhancer-promoter contacts in the partially extruded loop state. We propose a model of topological regulation whereby the transient, partially extruded states play active roles in transcription.

## Background

Topologically associated domains (TADs) and regulatory enhancer-promoter chromatin loops are largely formed by the cohesin complex through the process of CTCF-mediated loop extrusion^1,2^. Topological alterations and subsequent changes in enhancer-promoter (EP) contacts can modify gene expression^3,4^ and cause aberrant phenotypes^5–8^. CCCTC-binding factor (CTCF) can act as an extrusion barrier through its ability to bind and stabilize cohesin on DNA, serving to preferentially localize and anchor one or both ends of cohesin loops. Genes with promoter-proximal CTCF binding sites have been shown to have increased dependence on distal enhancers^9–11^, although the exact mechanisms involved are not well understood.

Although conventional 3C techniques give an impression of static loops, cohesin-mediated chromatin loops are actually dynamic with an extrusion rate of ~1kb/s^12^. Recent live cell-imaging studies of two TADs found that the fully extruded state with a loop formed between two convergent CTCF-bound anchors was present only 3-30% of the time^13,14^. While these findings suggest that CTCF loops spend the vast majority of their time partially-extruded, the partially-extruded state has not yet been well characterized.

Several studies have found evidence of promoter-proximal CTCF binding sites (CBS) having large impacts on EP contact frequencies and transcription^9–11^. Putting this together with the high prevalence of partially extruded CTCF-mediated loops, we hypothesize that promoter-proximal CTCF sites enable gene regulation by halting cohesin on one side while cohesin continues to extrude on the other side. Enhancers then slow down extrusion, thus enabling an increase in EP contacts without requiring a fully extruded loop. The relationship between EP contacts and transcription can be nonlinear such that small increases in EP contacts may cause large changes in transcription^3,4^. As a result, even minor decreases in extrusion rate through enhancer regions may affect gene expression.

The ability of MNase to efficiently digest naked DNA while sparing protein-bound DNA has been employed in various strategies to footprint the binding sites of proteins such as transcription factors with near base-pair resolution^15–18^. A key advantage of using MNase over sonication-based protocols is the shorter fragment size obtained, which directly leads to higher resolution TF binding site identification. More recently, MNase DNA fragmentation has also been applied to proximity ligation assays to map 3D genome architecture with nucleosome (~150 bp) resolution, enabling precise characterization of 3D architecture including at TAD boundaries and punctate enhancer-promoter interactions^19–22^. Since MNase HiChIP enables precise characterization of both TF-binding and 3D contacts, it is uniquely poised to define how CTCF enables 3D contacts.

To better characterize the partially extruded chromatin loop state, we first develop a computational technique for high-resolution footprinting of CTCF using MNase HiChIP data. We then employ this to study how, through its interaction with the looping factor cohesin, CTCF can facilitate long-range DNA contacts. We further characterize how the length of loops extruded by cohesin is affected by local chromatin state factors such as enhancer and RNA Pol II density.

## Results

### MNase HiChIP generates short, TF-protected and longer, histone-protected DNA fragments

We used Micrococcal nuclease (MNase) HiChIP^23^ with a CTCF antibody to profile 3D architecture in K562 cells, generating 150 bp reads with over 380 million unique pairwise contacts across four replicates. Briefly, following cell fixation with DSG and formaldehyde, chromatin is digested by MNase, immunoprecipitated to enrich for CTCF-bound DNA, and free ends are then ligated. After reverse-crosslinking, the resulting ligation products are sequenced from both ends and the mapping locations of the paired reads can be used to infer chromosomal locations of the physically interacting loci. In cases where the pre-ligation fragments are shorter than the read length it is also possible to infer the fragment length as the ligation junction position will be observed within one or both of the reads. If multiple fragments within a read are short enough to be aligned to distinct genomic locations, this is termed an ‘observed ligation’ (Fig. 1a, Supp Fig 1).

**Fig. 1.**
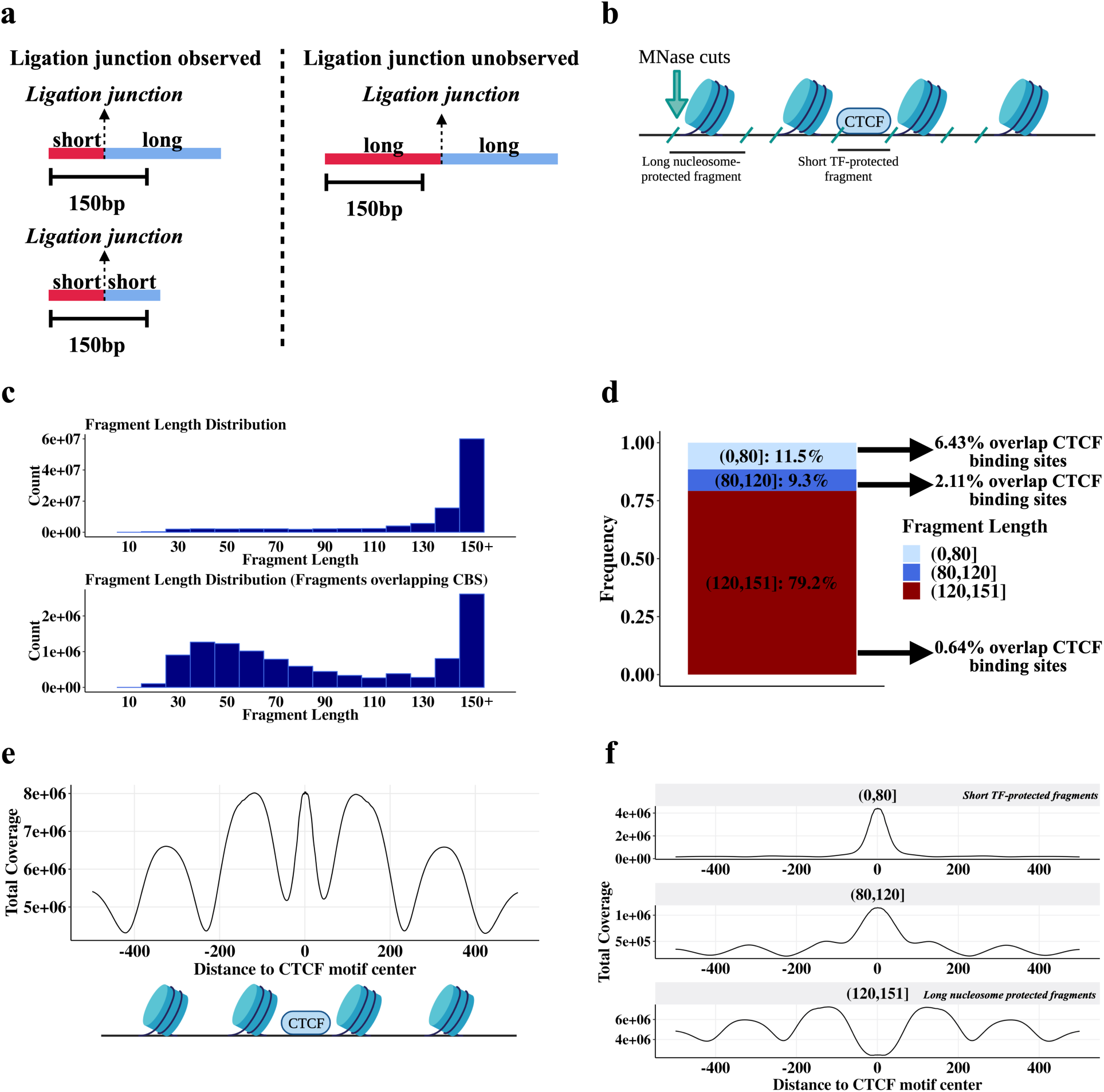
MNase CTCF HiChIP data contains short (~ <80 bp) CTCF-protected fragments and longer (~ >120 bp) nucleosome-protected fragments. **a** Schematic illustrating relationship between short fragments and observed ligations. **b** Schematic illustrating how the fragment length results from MNase cutting around bound proteins of different sizes. **c** Fragment length distribution for all fragments (top plot) and fragments overlapping occupied CTCF motifs (lower plot). Occupied CTCF motifs are defined here as CTCF motifs within 30 bp of a CTCF ChIP-seq peak summit. **d** Boxplot quantifying the frequency of different fragment lengths genome-wide and how often each fragment length group overlaps an occupied CTCF motif. Occupied CTCF motifs are defined here as CTCF motifs within 30 bp of a CTCF ChIP-seq peak summit. **e** Fragment coverage metaplot +/− 500 bp around CTCF binding sites. Schematic below the coverage metaplot illustrates the proteins producing these peaks. **f** Plot **(e)** stratified by fragment length.

As expected, due to the preference of MNase to selectively cleave DNA not shielded by bound proteins and the high abundance of histones in chromatin (Fig. 1b), the predominant fragment length is approximately 150 bp, indicative of cuts between nucleosomes^24^ (Fig 1c). We also noted a distribution of shorter fragment lengths, with 20% representing lengths shorter than 120 bp (Fig. 1d). A metaplot centered on CTCF binding site motifs shows an enrichment of 30-60 bp fragments suggesting that these shorter fragments represent CTCF-bound DNA (Fig. 1c)^2,25,26^. Consistent with this, we find that short (<80 bp) fragments have a 10-fold higher overlap frequency with CTCF motifs than long (>120 bp) fragments (Fig. 1d). This is similar to data from the MNase-based CUT&RUN assay that also results in short fragments protected by small proteins such as transcription factors^17^.

Fragment pileups at CTCF motif loci (Fig. 1e) show a strong enrichment of short fragments centered on the CTCF motif sequence, and a concomitant depletion of long fragments at motifs (Fig. 1f). Long fragments, in contrast, show peaks with a strong ~200 bp periodicity adjacent to the central CTCF binding site (Fig. 1f). This is consistent with the ability of CTCF to precisely position a series of nucleosomes adjacent to its binding site^25^. Note that while long (>120 bp) fragments are depleted at CTCF binding sites, they still represent a significant fraction of reads at these sites (Fig. 1c). This likely reflects that CTCF motif loci without a bound CTCF are frequently instead occupied by histones^25^, and even CTCF motifs with very strong CTCF ChIP-seq signal are not always occupied by a CTCF.

In summary, long fragments correspond to nucleosome-protected DNA whereas short fragments arise from TF-protected DNA. This is due to the different sizes of CTCF and histone octamers, with nucleosomes protecting about twice the amount of DNA that CTCF protects^25^. Since MNase cuts around bound proteins, the different protein sizes directly translate to different fragment lengths. Accordingly, we next filter out long, nucleosome-protected fragments and focus on short, TF-protected fragments to identify CBS.

### FactorFinder leverages the strand-specific bimodal distribution of short fragments around CBS to obtain precise detection of CTCF binding

In order to characterize CTCF-mediated chromatin loop interactions, we first set out to map CTCF loop anchors with high resolution. We take advantage of the difference in fragment lengths associated with CTCF-bound vs nucleosome-bound DNA to focus only on likely CTCF-bound fragments. Fragment lengths can be determined for all fragments with length less than 150 bp; the 150 bp read length results in censoring of fragments longer than 150 bp. While exact fragment lengths can be obtained for all fragments shorter than 150 bp, observed ligations require a shorter fragment length. This is because observed ligations require distinct mapping of fragments on either side of the ligation junction. Since at least ~25 bp are required to align a sequence to the reference genome, this results in fragments characterized as observed ligations having a maximum fragment length of ~125 bp, sufficient for the identification of most CTCF-protected DNA fragments. Consequently, the fraction of informative, CTCF-protected fragments decreases with shorter sequencing read length (Supp Fig 1). The effect of subsetting the CTCF HiChIP dataset to only short fragments (<125 bp, identified by the proxy of an observed ligation), is shown in Fig 2a,b. These shorter, presumably CTCF-protected fragments, are overwhelmingly located immediately adjacent to CTCF motifs.

**Fig. 2.**
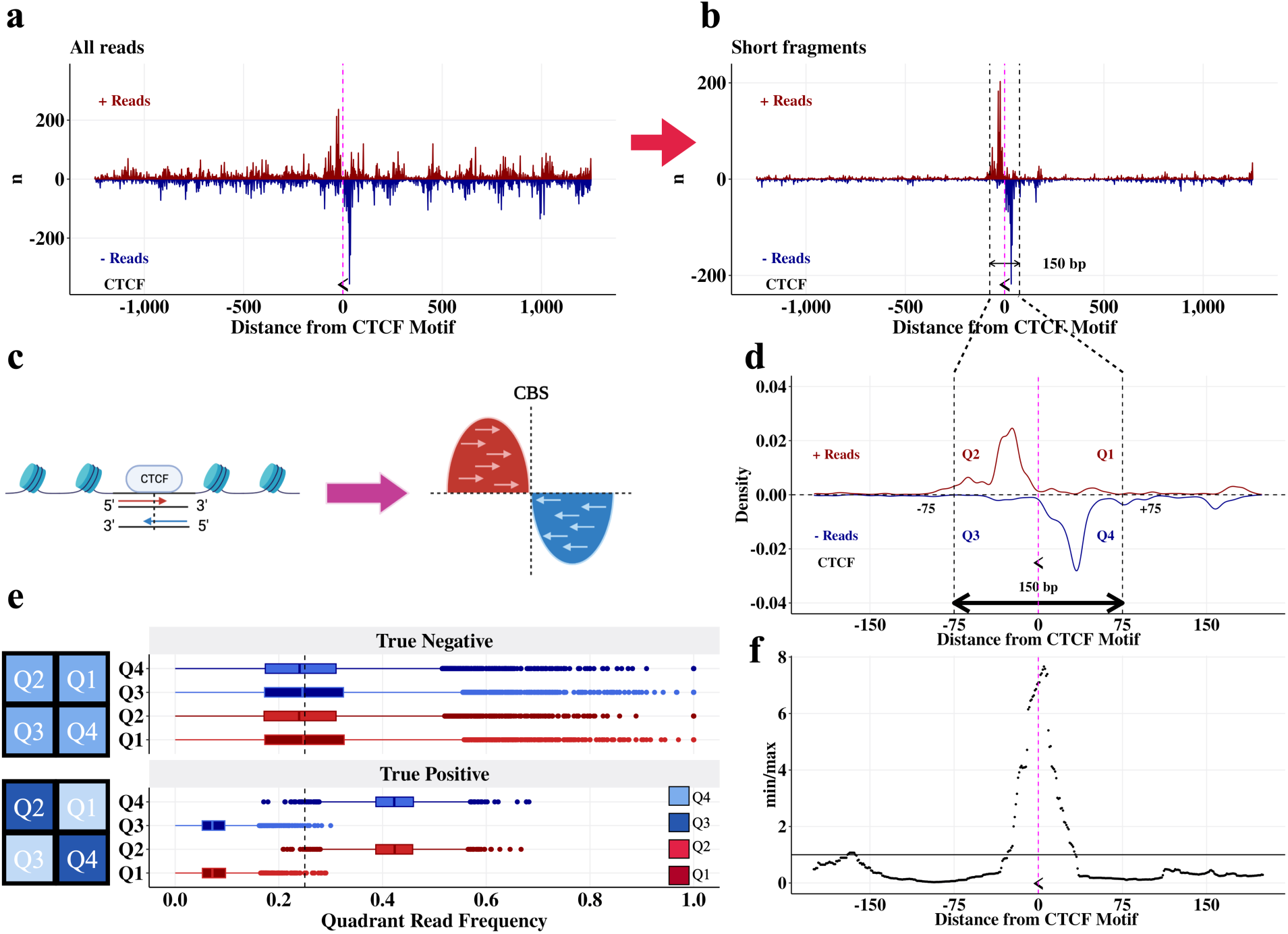
True CTCF binding sites have a bimodal strand-specific distribution centered on the CTCF motif. **a** Unfiltered reads +/− 1250 bp around a CTCF binding site located on the negative strand (chr1: 30,779,763 - 30,779,781). The midpoint of the CTCF motif is marked with the symbol “<”, representing that it is on the negative strand, and a pink line. **b** Plot **(a)** filtered to observed ligations (equivalently, short fragments.) **c** Schematic demonstrating the bimodal read pile-up around a CTCF binding site. **d** Plot **(b)** as a density plot and zoomed in on the CTCF motif, with quadrant annotations. **e** Distributions of reads in quadrants for true negative and true positive CTCF binding sites in DNA loop anchors. True positives are defined as CTCF motifs that are the only CTCF motif in a loop anchor and within 30 bp of a CTCF ChIP-seq peak. True negatives are areas of the loop anchors with one CTCF motif that are at least 200 bp from the CTCF motif. Schematics of the quadrant read pile-up patterns are shown next to the corresponding true positive and true negative boxplots. **f** *FactorFinder* statistic 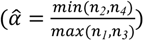 for plot **(d)** peaks at the CTCF motif.

Sequencing of short, CTCF-protected fragments results in a bimodal read distribution centered on the CBS, with read 5’ location peaks observed upstream (positive strand) and downstream (negative strand) of the CBS (Fig. 2c). We refer to these regions as quadrants 2 and 4 (Q2 and Q4) respectively (Fig. 2d, e). In contrast, reads from the positive strand downstream of the CBS (Q1) and negative strand upstream of the CBS (Q3) correspond to fragments with MNase cut sites underneath CTCF-protected DNA, and therefore reflect a lack of CTCF occupancy. CTCF binding therefore produces an enrichment of reads in Q2,Q4 and a depletion of reads in Q1,Q3 (Fig. 2e). At sites without protein binding, MNase can cut at any location resulting in no enrichment of reads in Q2 and Q4 compared to Q1 and Q3 (Fig. 2e). As a result, we can determine CTCF binding by testing if there are significantly more reads in Q2 and Q4 than Q1 and Q3 (Fig. 2f).

We can consider each read as an independent draw from a multinomial distribution with four categories corresponding to the four quadrants. Under the null hypothesis, each read has equal probability of belonging to any of the four quadrants *Q_i_*, *i* ∈ {*1,2,3,4*}. Because true CTCF binding induces a strong read pile-up in *<u>both</u>* quadrants 2 and 4 in addition to a depletion of reads in quadrants 1 and 3 (Fig. 2d, e, f), we test for an enrichment of reads in Q2 and Q4 compared to Q1 and Q3 by estimating the *FactorFinder* statistic 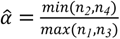, where *n_i_* is the number of reads in *Q*i. We then test if 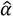 is significantly greater than 1. Note that min and max are used to enforce that both quadrants 2 and 4 must have more reads than both quadrants 1 and 3; using the average would enable read pile-ups that occur in quadrant 2 or 4 (but not both) to be spuriously called as CTCF binding events.

To evaluate the significance of 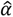 at a particular total read count 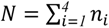, we simulated 100 million samples under the null hypothesis that each fragment is equally likely to occur in any of the four quadrants. This was done at each total read count ranging from 5 to 500. P-values at read counts beyond 500 are very similar to those at 500, so 500+ read counts are treated as bins with 500 total read count (Supp Fig 2). The empirical CDF of the 100 million 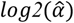 at a given total read count was then computed and used to evaluate the probability of observing a value more extreme than 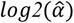 under the null hypothesis. The empirical CDF was evaluated at a sequence of possible 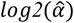 between 0 and 5 at step sizes of 0.01(this corresponds to 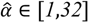.) This approach produces the same p-values as using 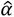 instead of 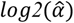, but using the log enables smaller step size at large values of 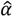. After acquiring the grid of p-values for each 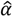 at a given read count *N*, we match the observed 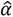 at a read count of *N* with the corresponding p-value from the table. Because this approach only requires quadrant-specific read counts to match with the given table of p-values, it is very computationally efficient. Furthermore, by using the multinomial framework we place no assumptions on the reads within each quadrant being distributed as poisson, negative binomial, or another distribution. The only assumption we make is that in the event of no CTCF binding, the reads are equally distributed amongst the four quadrants. We have shown this assumption holds in Figures 2c, d, e.

In brief, we have shown that short fragments exhibit a strand-specific, bimodal distribution centered on the CBS. This distribution arises from MNase cutting around a bound CTCF and subsequent sequencing 5’ to 3’ of the DNA. Significance is assessed through a multinomial framework, which has the advantage of not placing any assumptions on the distribution of reads within each quadrant. Now that we have explored the theory behind *FactorFinder*, we demonstrate its ability to identify CBS with high resolution and accuracy.

### Model evaluation

*FactorFinder* uses a biologically-informed model that takes advantage of the distribution of short fragments around a CTCF binding site to pinpoint CTCF binding. Additionally, our use of a multinomial framework for significance evaluation avoids placing any distributional assumptions on the reads within a quadrant. We then sought to benchmark our CTCF binding site identification performance using CTCF motif locations^27^, CTCF ChIP-seq peaks^28^, and loop anchors identified by FitHiChIP at 2.5kb resolution^29^.

We define a high stringency true positive set of CTCF binding sites as CTCF motifs in loop anchors that are located within 30 bp of a CTCF ChIP-seq peak summit. To avoid ambiguity due to multiple closely spaced motifs, we further selected only those motifs that are unique within a 2.5kb loop anchor. Using this true positive set, we observe that the *FactorFinder* statistic, 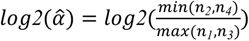 has signal greater than 0 (equivalently, 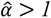) almost exclusively within 20 bp of the CTCF motif center and centered on 0 bp from the CTCF motif center (Fig. 3a). Using this same set of true positive sites (false negatives are the regions of the loop anchors >200 bp from a CTCF motif), we achieve > 90% precision and > 90% recall at a p-value threshold of 1e-05, and maintain high recall and precision at all p-value thresholds < 1e-05 (Fig. 3b). This high level of recall and precision is achieved because of the very different *FactorFinder* statistic distributions for true positives and true negatives (Fig. 3c).

**Fig. 3.**
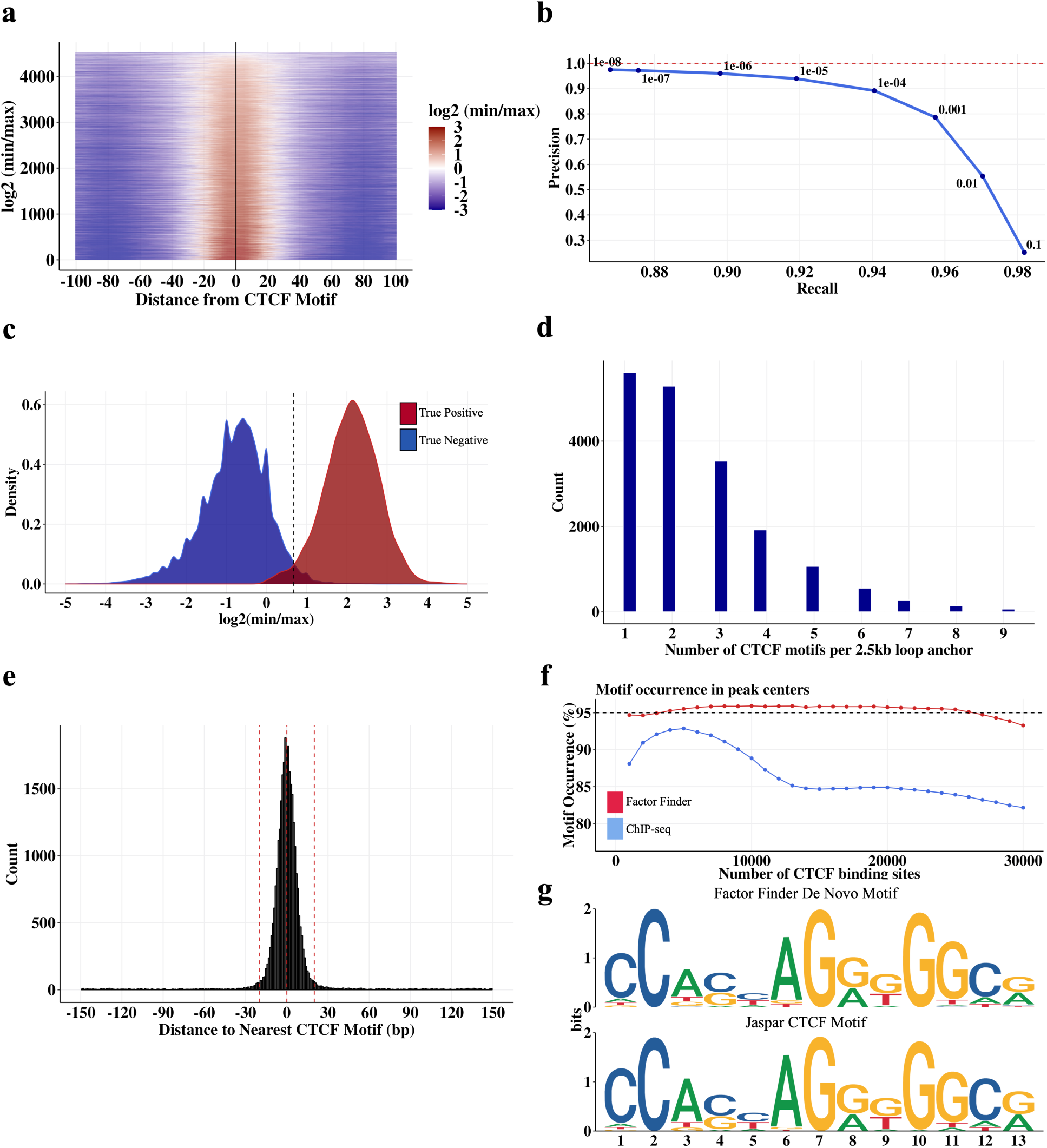
CTCF binding sites identified by *FactorFinder* with single basepair resolution in MNase K562 CTCF HiChIP data. **a** Heatmap of log2(min/max) as a function of distance between *FactorFinder* peak center and CTCF motif center within loop anchors. Only CTCF motifs that are unique within a loop anchor and within 30 bp of a CTCF ChIP-seq peak are used. **b** Precision recall curve for true negative and true positive CTCF binding sites in DNA loop anchors. True positives are defined as in **(a)**. True negatives are areas of the loop anchors in **(a)** that are at least 200 bp from the one CTCF motif. Precision is calculated as TP / (TP + FP), recall is calculated as TP / (TP + FN). **c** *FactorFinder* statistic density plots using the same set of true positives and true negatives as **(b). d** Distribution of the number of CTCF motifs in a 2.5kb loop anchor. **e** Histogram with 1 bp bin size depicting *FactorFinder* resolution for all peaks genome-wide (not just in loop anchors). **f** Motif occurrence in ChIP-seq and *FactorFinder* peak centers genome-wide. Motif occurrence is calculated as % peak centers within 20 bp of CTCF motif. Only peak centers within 150 bp of a CTCF motif are used for this figure. **g** 30 bp sequences centered on genome-wide *FactorFinder* peak centers produce a de novo motif (top) that matches the core JASPAR CTCF motif (bottom).

Because 70% of loop anchors defined with 2500 bp resolution contain multiple CTCF motifs (Fig. 3d), higher levels of precision are often needed to determine the specific CTCF motif(s) mediating a CTCF loop. Examining the effectiveness of *FactorFinder* genome-wide, we observe that almost all *FactorFinder* peak summits (93%) are within 20 bp of a CTCF motif center, with a median separation of 5 bp (Fig. 3e). Quantifying accuracy using motif occurrence within 20 bp of a peak summit, we find that *FactorFinder* maintains ~95% motif occurrence while ChIP-seq declines to less than 85% motif occurrence (Fig. 3f). Applying the motif discovery tool STREME^30^ to 30 bp sequences centered on the *FactorFinder* peak summit produces a motif sequence that exactly matches the core JASPAR CTCF motif (Fig. 3g), further supporting *FactorFinder*’s ability to identify true CTCF binding sites.

### CTCF and Cohesin occupancy footprints

We next examined the length characteristics of MNase HiChIP fragments overlapping individual CTCF motifs, to infer the presence and identity of the protein occupying the locus. For motifs with non-zero coverage, we observed long, 150+ bp fragments, as shown for three representative motifs in Figure 4a. These fragments likely represent cells with a nucleosome located at the motif locus, and are observed at CTCF motifs genome-wide (Fig. 1c). In addition, for a large subset of CTCF motifs, we also observed short, sub-nucleosome sized (<115 bp) fragments (Fig. 4a, Fig. 1c), likely instead representing DNA protected by CTCF.

**Fig. 4.**
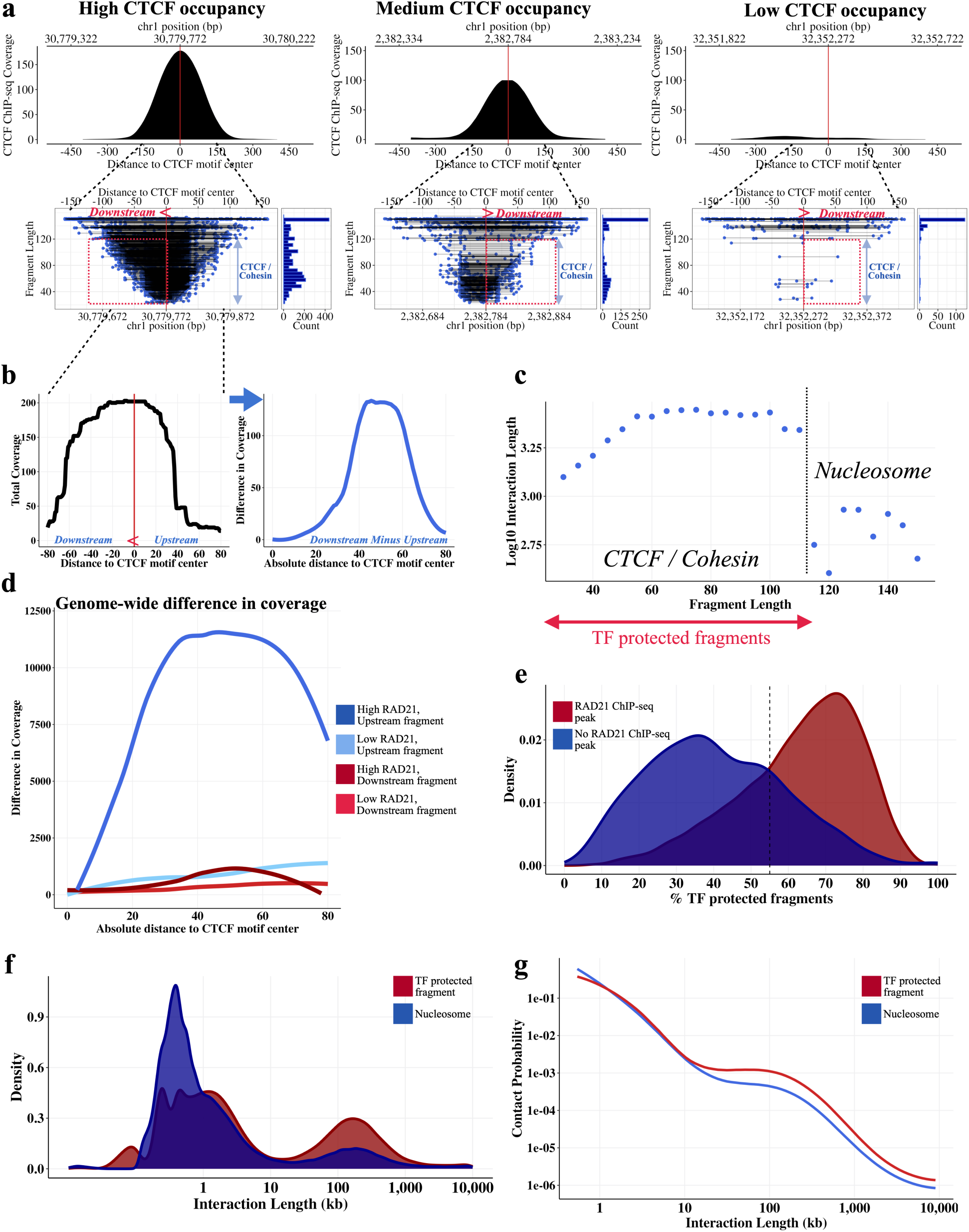
Cohesin and CTCF-protected fragments identified in CTCF MNase HiChIP. **a** High, medium, and low CTCF occupied motifs. Cohesin footprint is observed downstream of the CBS for high and medium CTCF occupancy motifs. For each occupancy level, CTCF ChIP-seq (top) and all fragments overlapping the CTCF motif (bottom left) are depicted, along with the corresponding fragment length histogram (bottom right). **b** Locus-specific high CTCF occupancy figure from **(a)** as a coverage plot (left figure), difference in coverage between downstream and upstream coverage (right figure). **c** Plotting median log10 interaction length as a function of fragment length suggests presence of nucleosome vs TF-protected fragments. Only left fragments overlapping CTCF (+) motifs with start and end at least 15 bp from the CTCF motif were included in this graph to remove confounding by MNase cut site. Using this figure, we are approximating CTCF +/− cohesin-protected fragments as those with fragment length < 115, start and end at least 15 bp from the motif center. **d** Difference in coverage (downstream - upstream) across all CBS shows an increase in coverage downstream of the CTCF motif for upstream fragments underlying CBS with a strong adjacent RAD21 ChIP-seq peak. **e** CTCF motifs that have a nearby RAD21 ChIP-seq peak (within 50 bp) have a larger proportion of TF-protected fragments. **f** TF-protected fragments have a noticeably larger bump in density of long range interactions compared to nucleosome-protected fragments. Fragments were first filtered to those with start and end at least 15 bp from the motif. TF-protected fragments were then defined as fragments with length < 115 bp while nucleosome-protected fragments are fragments with length at least 115 bp. **g** P(S) curve for fragments depicted in **(f)**.

A closer examination of the TF-scale fragments at *FactorFinder*-identified bound motifs reveals that they tend to exhibit a skew towards the downstream side of the CTCF motif (Fig. 4a, b, c), suggesting a preferred location for the protein(s) protecting the region from MNase cleavage. We considered cohesin as a potential candidate, given a recent finding that cohesin is stabilized on DNA through a specific interaction with the N terminus of the CTCF protein^2^, which localizes to the downstream side of the CTCF binding site.

Given CTCF’s role in mediating DNA looping we investigated whether the CTCF-adjacent protected footprint might relate to 3D architecture within the cell. We used HiChIP pairwise interaction data where each ligation event reflects a single-cell point-to-point contact, to classify each CTCF motif-overlapping fragment as either ‘upstream” or ‘downstream’, depending on its relationship to its interaction partner. Upstream fragments have long range contacts downstream of the motif, and therefore have looping contacts in the same direction as a chromatin loop mediated by cohesin bound to the N terminus of the CTCF protein. Examining the difference in coverage downstream and upstream of CBS genome-wide, we observe that upstream fragments overlapping CBS with an adjacent strong RAD21 ChIP-seq peak have substantially more adjacent coverage in the ~60 bp region downstream compared to upstream of the motif, while downstream fragments and CBS with weak adjacent RAD21 ChIP-seq peaks exhibit no difference (Fig. 4d). This finding further suggests that the CTCF-adjacent factor is associated with loop formation.

To further investigate whether the TF footprints identified at CTCF motifs might relate to an architectural role, we used HiChIP data to characterize their interaction patterns. We found that TF-protected fragments (<115 bp) had contacts at substantially longer genomic distances than nucleosome-protected fragments (Fig. 4c), suggesting that the TF presence may facilitate long range interactions. Furthermore, we computed the frequency of TF-protected fragments at all *FactorFinder*-identified CTCF bound sites, and found that it is strongly associated with the presence of a RAD21 ChIP-Seq peak at the motif ^28^ (Fig 4e).

Examination of the interaction length distribution shows that, as expected, the majority of interactions occur within a linear separation of less than 10kb. The fraction of long-range (>10kb) interactions, however, is significantly enriched (3.5-fold, p < 10^-10^) for short TF-protected fragments as would be expected if these footprints represent CTCF/cohesin (Fig. 4f). Similarly, an examination of the P(s) curve, showing contact probability as a function of linear distance, reveals a decreased attenuation in contact probability at longer interaction lengths (Fig. 4g). Taken together, these findings suggest that we can classify CTCF HiChIP interaction data based on footprint/fragment size as involving either unoccupied CTCF sites that tend to have short-range chromatin interactions, or CTCF/cohesin occupied sites that, presumably through loop extrusion, are able to make long-range contacts.

### Active enhancers and gene transcription hinder cohesin-mediated loop extrusion

Using the techniques described above, MNase HiChIP enables us to simultaneously locate CBS at high resolution, identify footprints of bound proteins, and interrogate specific chromatin contacts at the single molecule level. We next sought to leverage these data to characterize cohesin extrusion dynamics in a range of genomic contexts.

We first estimated the frequency of fully extruded CTCF-CTCF chromatin loops genome-wide. By obtaining fragments overlapping CTCF binding sites and estimating the fraction of interaction partners overlapping a downstream convergent CTCF motif, we obtain 5% as the genome-wide frequency of the fully extruded CTCF-CTCF state.. We also find a wide CBS to CBS variability with an estimated range of ~1-10% (Fig. 5a). This suggests that most CTCF-anchored chromatin contacts at the single-cell level are in the ‘extruding’ state, rather than joining two CTCF sites. These ranges are consistent with two recent locus-specific live cell imaging studies, which found that the fully extruded loop state is rare at the *Fbn2* TAD^13^ and an engineered TAD on chr15^14^, occurring ~3-6%^13^ and ~20-30% of the time^14^ respectively. Note that the 20-30% estimate corresponds to a loop existing between any combination of three CBS (+) and three CBS (−).

**Fig. 5.**
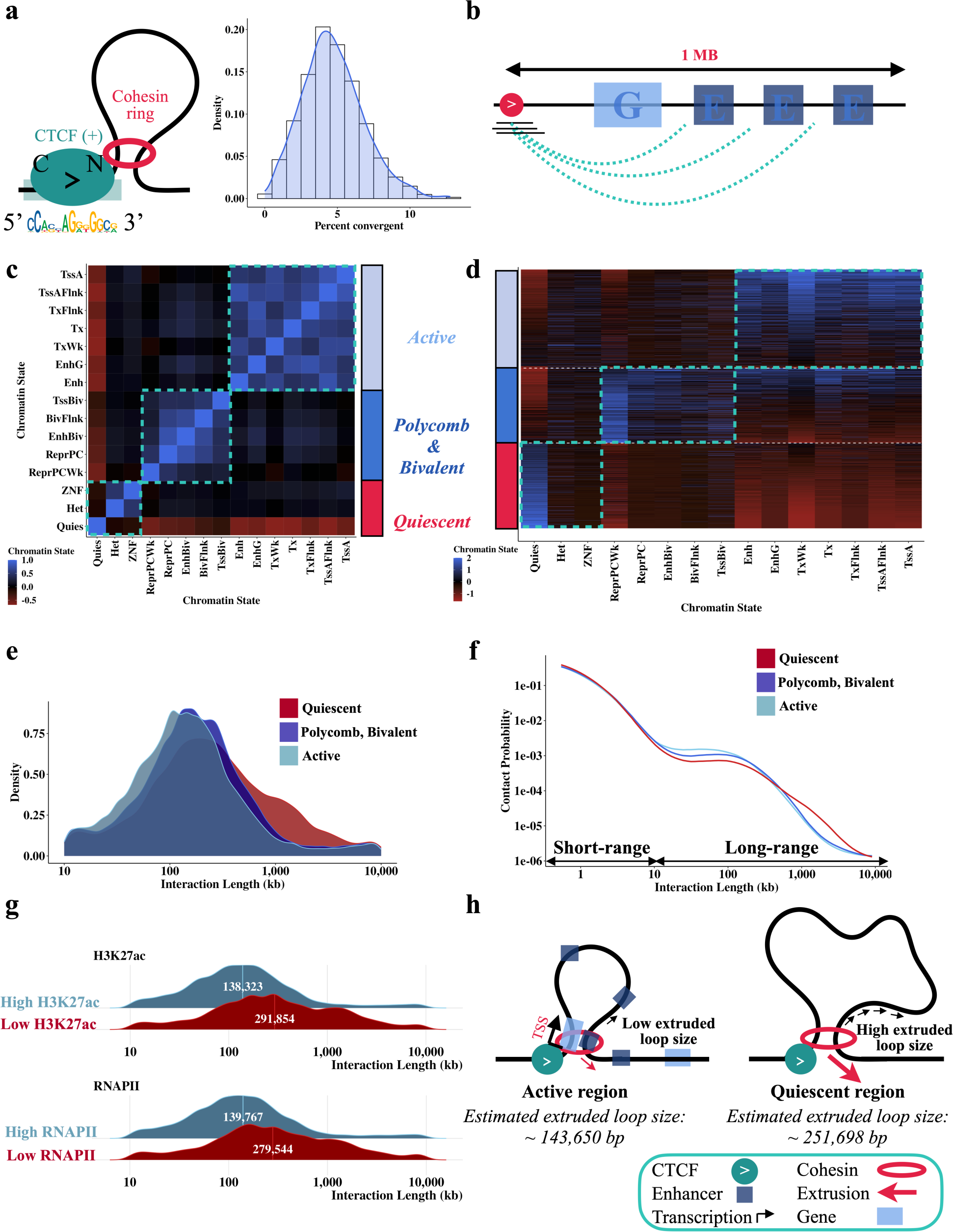
Cohesin extrudes further through quiescent regions than active regions. **a** Most CTCF-mediated looping contacts do not reflect the fully extruded state. Estimate is obtained using left TF-protected (start and end at least 15 bp from motif center, length < 115) fragments that overlap *FactorFinder* identified CBS (+) and have an interaction length greater than 10kb. For each CBS with at least 50 long-range TF-protected fragments overlapping the motif, % convergent is calculated as the number of interaction partners overlapping CTCF (−) motifs / total number of fragments at motif. Because this estimate is conditional on CTCF binding at the anchor, we divide estimates by two to account for the ~50% occupancy of CTCF^34^. **b** Depiction of how regions were annotated using ChromHMM. Correlation **(c)** and fragment **(d)** heatmaps for ChromHMM annotated unique 1 MB regions downstream of left fragments overlapping CTCF (+) binding sites. All other plots in this figure are filtered to TF-protected (fragment length < 115 bp, start and end at least 15 bp from motif center) fragments. Density **(e)** and P(S) curves **(f)** for chromatin state clusters shown in **(c,d)**, filtered to the top 20%. Chromatin annotations making up each cluster are added together and quantiles are obtained to determine fragments in the top 20% of active chromatin, quiescent chromatin, and bivalent / polycomb chromatin. **g** Ridge plots for the bottom 10% quantile (“Low”) and top 10% quantile (“High”) of H3K27ac bp and number of RNAPII binding sites. ChIP-seq from ENCODE was used to annotate 1 MB downstream of left fragments overlapping CBS (+) for this figure. **h** Diagram illustrating differences in extrusion rates between active and quiescent chromatin states, with numbers obtained from Supp Fig. 3.

We next sought to use our data to examine how cohesin extrusion is impacted by chromatin context. Since HiChIP libraries are a snapshot of millions of cells, we can estimate dynamic extrusion parameters (primarily the average loop size extruded by cohesin^31^) from the interaction length distribution. To determine the impact of chromatin state on cohesin extrusion, we first annotated the 1 MB regions downstream of *FactorFinder* identified CBS with ChromHMM states^32^ (Fig. 5b) to characterize the DNA through which a cohesin anchored at the CBS would extrude through. Due to the highly correlated nature of ChromHMM annotations (Fig. 5c, d), we then divided the genome into three main chromatin state categories to uniquely classify each 1 MB region as either active, polycomb/bivalent or quiescent (Fig. 5d). CTCF/cohesin-protected fragments overlapping CBS were accordingly annotated with the corresponding motif-level chromatin state group, and extruded loop size estimates were obtained for each chromatin state based on the fragment-level interaction lengths.

Interestingly, we find that cohesin extrudes 1.75 times further through quiescent regions (252kb) than through active regions (144kb), corresponding to a difference in average extruded loop size of ~110kb, p < 10^-10^ (Fig. 5e, Supp Fig. 3, Supp Fig. 4 right). The P(s) curve, a plot of interaction decay with distance, confirms a depletion of the longest-range interactions in active regions (Fig 5f). This estimate for quiescent regions is consistent with a live cell imaging study of the *Fbn2* locus in the absence of transcription that estimated a processivity of 300kb^13^. As quiescent regions are characterized by low TF binding, low transcription, and minimal histone modifications^33^, we hypothesized that the substantial difference in extruded loop size relates to gene activity and enhancer density obstructing loop extrusion. Consistent with this, we found that higher levels of H3K27ac and RNA Pol II binding in the 1MB region downstream of the CBS strongly correlate with lower average extruded loop size (Fig. 5g).

We sought to establish that the observed differences in loop extrusion length as a function of chromatin state are not confounded by locus-specific effects on cohesin extrusion. Each CBS has locus-specific genetic architecture and a different number of overlapping fragments, so we fit a linear mixed effects model to account for this group-level heterogeneity. Specifically, we compute the ‘cohesin effect’ on loop length, defined as the increase in average interaction length for CTCF/cohesin bound fragments compared to nucleosome bound fragments for each CBS. Controlling for the background interaction frequency of a region in this way confirms that cohesin-associated loops are significantly shorter in active chromatin (Supp Fig. 4 left). Taken together, these findings imply that gene and enhancer activity impede cohesin translocation (Fig. 5h).

## Discussion

We have developed *FactorFinder*, a transcription factor footprinting method for MNase HiChIP data and used it to identify CTCF binding sites with near base-pair resolution. We show that the DNA protection footprints of nucleosomes and transcription factors can be readily distinguished based on pre-ligation fragment size and strand origin and use these features to identify CTCF binding sites. Significance is then assessed through a multinomial approach, which avoids placing distributional assumptions on read counts. Using this method, the median distance between *FactorFinder* peak summits and motif center is 5 bp, with 93% of peak summits identified within 20 bp of a CTCF motif center.

We then leverage this methodological advance to investigate how chromatin state affects cohesin extrusion dynamics. A close examination of CTCF-protected fragments revealed an additional CTCF-adjacent footprint downstream of the CBS, which we propose represents cohesin given its positioning relative to looping orientation as well as its strong association with both long range interactions and cohesin occupancy. We estimated the frequency with which a CTCF bound locus forms a loop with a downstream CTCF site and found that it varies considerably from CBS to CBS, with a genome-wide range from ~1-10%. This is consistent with recent live-cell imaging work that found that CTCF-mediated loops predominantly exist in the partially extruded state at two studied loci^13,14^.

We next sought to characterize how cohesin impacts genome contacts in different chromatin contexts. To this end, we employed our high-resolution *FactorFinder* identified CBS and HiChIP 3D contact information to look at differences in extruded loop size in regions with different chromatin states. We observe an approximately 2-fold increase in extruded loop size comparing quiescent chromatin to active chromatin, and this effect is similarly observed when examining the impact of H3K27ac and RNAPII binding. Our finding that RNAPII binding obstructs cohesin-mediated loop extrusion is consistent with two recent studies that investigated RNAPII’s impact on cohesin through RNAPII and enhancer perturbations^35^ as well as polymer simulations, CTCF depletion, and Wapl knockout experiments^36^. These substantial differences in average extruded loop size observed for different levels of RNAPII binding and H3K27ac suggest that gene and enhancer activity obstruct cohesin-mediated loop extrusion.

The obstruction of cohesin by gene and enhancer activity implies a model of CTCF-mediated gene regulation where a fully extruded, stable, and convergent CTCF-CTCF loop is not required for CTCF to mediate enhancer-promoter contacts. Instead, a promoter-proximal CTCF can halt cohesin next to the TSS of a gene while cohesin continues to extrude on the other side, effectively behaving as an enhancer recruiter. Cohesin slowing down through enhancer regions would then enable an enrichment of enhancer-promoter contacts without requiring a stable CTCF-CTCF loop (Fig. 6). This attenuation in cohesin extrusion may also provide a mechanism relating gene regulation to the presence of RNAPII at enhancers^37^.

**Fig. 6.**
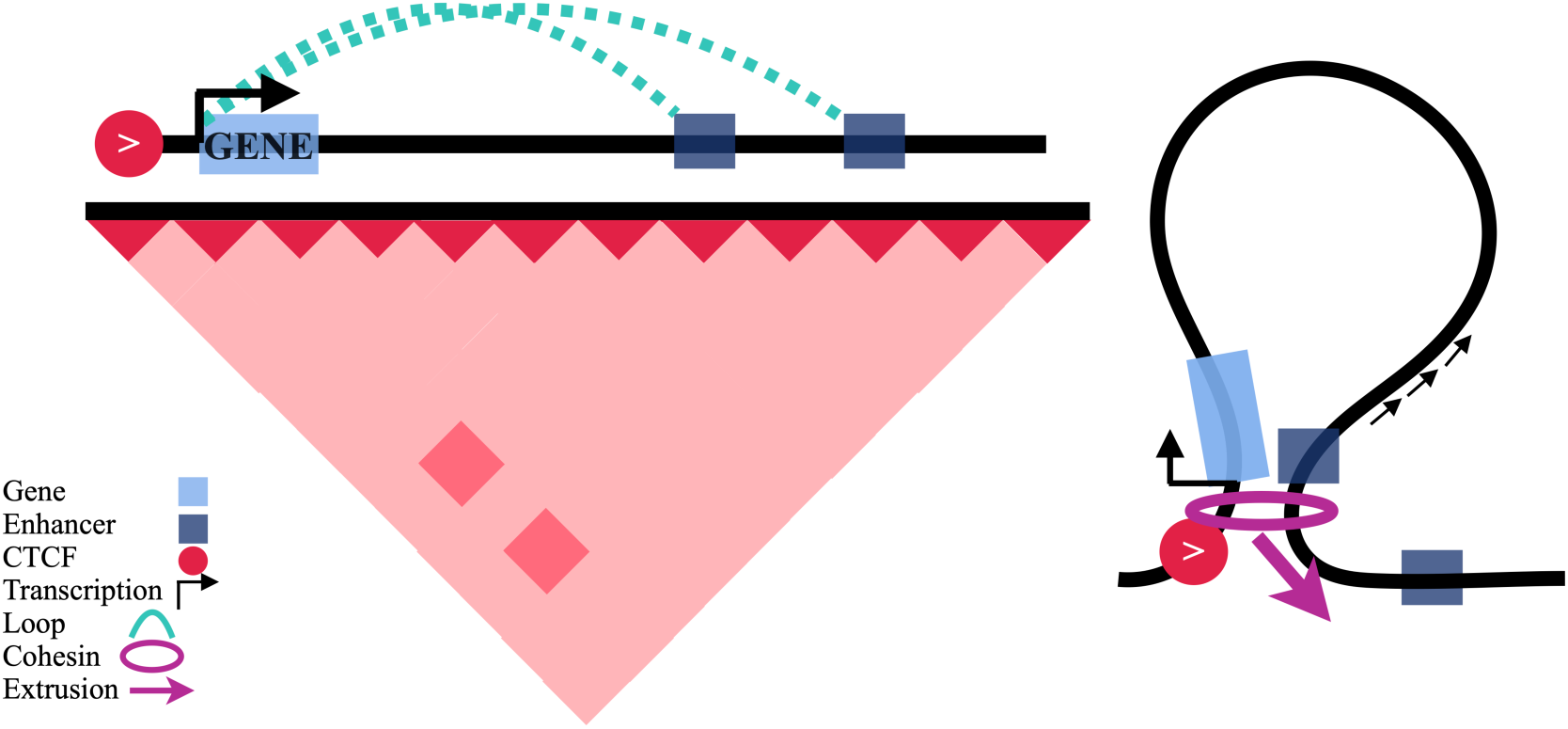
Schematic of proposed model whereby single promoter-proximal CTCF sites enable an enrichment of enhancer-promoter contacts.

The dynamic CTCF-mediated enhancer-promoter contact model proposed here is consistent with recent findings that promoter proximal CTCFs have important roles in gene regulation^9–11^, that enhancer-promoter contacts are unstable^38,39^, and that CTCF and cohesin-mediated chromatin loops are dynamic^13,14^. The dynamic nature of EP contacts has contributed to the development of the “kiss and kick” model^40^ as a potential explanation for how enhancers and promoters come into contact but move away from each other at the time of transcription. Our findings are compatible with the “kiss and kick” model, but additionally suggest a potential mechanism by which distal enhancers can locate gene promoters without being stuck in a stable conformation. This model would use promoter- or enhancer-proximal CTCF sites to enable distal enhancers to both come into contact with gene promoters and subsequently disengage during transcription. In this way, CTCF’s role in long-range enhancer promoter contact would be as a dynamic functional element recruiter instead of mediating continual stable contact between distal enhancers and gene promoters.

## Materials and methods

### CTCF MNase HiChIP

Four MNase K562 CTCF HiChIP (150 bp paired-end) libraries were generated using the Cantata Bio / Dovetail Genomics MNase HiChIP kit. CTCF MNase HiChIP was performed as described in the Dovetail HiChIP MNase Kit protocol v.2.0. Briefly, 5 million K562 cells per sample were crosslinked with 3mM DSG and 1% formaldehyde and digested with 1ul MNase (“YET” samples) or 2ul MNase (“GW” samples) in 100ul of 1X nuclease digestion buffer. Cells were lysed with 1X RIPA containing 0.1% SDS, and CTCF ChIP was performed using 1500ng of chromatin (40-70% mononucleosomes) and 500 ng of CTCF antibody (Cell Signaling, cat #: 3418). Protein A/G beads pull-down, proximity ligation, and library preparation were done according to the protocol. Libraries were sequenced to a read depth of ~172 million paired end reads per sample on the Illumina Nextseq 2000 platform.

### Software implementation

Preprocessing, analysis and figure code used in this paper are available at https://github.com/aryeelab/cohesin_extrusion_reproducibility. Data figures in this paper were made in R v.4.1.2 using ggplot.

### Data availability

Raw and Processed HiChIP data produced in this study will be uploaded to NCBI GEO (GSE Record ID pending).

K562 ChIP-seq RAD21 BED file (Accession ID: ENCFF330SHG), CTCF BED file (Accession ID: ENCFF736NYC), CTCF bigWig signal value (Accession ID: ENCFF168IFW), RNAPII BED file (Accession ID: ENCFF355MNE), and H3K27ac BED file (Accession ID: ENCFF544LXB) were obtained from ENCODE, and CTCF motifs were obtained from the R package *CTCF* ^27^ (annotation record: AH104729, documentation: https://bioconductor.org/packages/release/data/annotation/vignettes/CTCF/inst/doc/CTCF.html).

## Methods

### Data Processing

4 replicates of K562 MNase CTCF HiChIP data were aligned to the reference genome using the BWA-MEM algorithm^41^. Ligation events were then recorded using pairtools parse v. 0.3.0^42^, PCR duplicates were removed, and the final pairs and bam files were generated. HiChIP loop calls were then made using FitHiChIP Peak to Peak^29^ with 2.5kb loop anchor bin size. The MNase HiChIP processing protocol is based on guidelines from https://hichip.readthedocs.io/en/latest/before_you_begin.html. Reproducible code is available at https://github.com/aryeelab/cohesin_extrusion_reproducibility.

### Identification of significant motifs

We use CTCF motifs identified as significant (p < 1e-05) by *FactorFinder* as the set of CTCF binding sites. This p-value threshold was chosen based on the precision recall curve (Fig. 3b), and corresponds to a maximum FDR q-value of 3e-04.

### Multiple Testing

For genome-wide footprinting analysis adjustment for multiple testing, CTCF motifs are assigned the p-value of the closest *FactorFinder* sliding window. The Benjamini-Hochberg method^43^ was used to obtain q-values.

### Estimating cohesin footprints

The cohesin footprint is observed by obtaining motif-level coverage estimates +/− 80 bp around CBS, summing up the coverage across all motifs (within strata), and subtracting the upstream coverage from the downstream (downstream coverage - upstream coverage) at each base pair. Note that downstream and upstream are defined relative to the motif strand, so downstream is to the “left” of CBS (−) and to the “right” of CBS (+) in terms of reference genome base pairs. The aforementioned strata are defined by RAD21 ChIP-seq signal level (high vs low) and whether the fragment is the upstream or downstream interaction partner in its pair. RAD21 ChIP-seq high and low correspond to the top 25% and bottom 25% of ChIP-seq signal value of the adjacent (within 50 bp of CBS) RAD21 ChIP-seq peak. Note that only mid-size (fragment length between 80 and 120), long range fragments (interaction length > 10kb) are used for this analysis.

### Estimating the fully extruded state

We estimated a genome-wide range for the fully extruded state by obtaining CTCF/cohesin-protected upstream fragments overlapping CBS (+) and estimating the fraction of interaction partners overlapping a downstream convergent negative strand CTCF motif. CBS (+) were required to have at least 50 CTCF/cohesin-protected upstream fragments overlapping the motif to enable sufficient sample size for the motif-specific percent convergent calculation. We then accounted for CTCF occupancy (estimated as ~50%)^34^ by dividing this estimate by two. The point estimate (5%) is the number of interaction partners overlapping a downstream convergent negative strand CTCF motif genome-wide / the total number of fragments genome-wide, and the range (1-10%) are the 1st and 99th percentile of the CBS-level CTCF-CTCF chromatin loop estimate.

### Determining extruded loop size as a function of chromatin state

We used upstream fragments overlapping CTCF binding sites (+) for this analysis. 1 MB regions downstream of the CBS (+) were annotated using ChromHMM^32^ to quantify the percentage of bp assigned to each of the 15 chromatin states. To simplify annotation, we grouped the 15 chromatin states into three categories (quiescent, polycomb/bivalent, and active) based on their correlation (Fig 5c). Regions were clustered using Ward’s hierarchical clustering method^44^ (Fig 5d.). For extrusion dynamics analyses (Fig 5e,f,h), each of the three chromatin categories was represented by the 20% of regions with the highest fraction of DNA in this state. Extruded loop size was then estimated as the average log10 interaction length for each annotation. Only long range TF-protected fragments (start and end at least 15 bp from the motif center, length < 115, interaction length > 10kb) were included in this estimate.

Similarly, high/low H3K27ac corresponds to the top 10% and bottom 10% of the number of basepairs covered by H3K27ac ChIP-seq peaks in the 1 MB regions downstream of CBS (+). High/low RNAPII corresponds to the top 10% and bottom 10% of the number of RNAPII ChIP-seq peaks located in the 1 MB regions downstream of CBS (+). Extruded loop size estimates were obtained in the same way for these annotated regions; long range TF-protected fragments were used to estimate the average log10 interaction length.

### Directionality of CBS-adjacent nucleosome position signal

Interestingly, the strength of the nucleosome positioning signal is related to the orientation of the DNA contact. Stratifying nucleosome-bound fragments based on whether they are the upstream or downstream long-range (>10kb) fragment in a pair (effectively single-cell left or right loop anchor) produces a differential nucleosome signal inside and outside the loop (Supp Fig. 5). For both upstream and downstream nucleosome-bound fragments, the nucleosome closest to the CTCF binding site and inside the loop exhibits a substantially stronger signal than the closest nucleosome outside the loop. HiChIP ligations are unlikely to fully account for this signal as a previous study using MNase-seq also showed a directional nucleosome preference around CBS (see Fig. 1a), although this result was not noted in the text^25^.

## Supplementary Figures

**Supplementary Figure 1.**
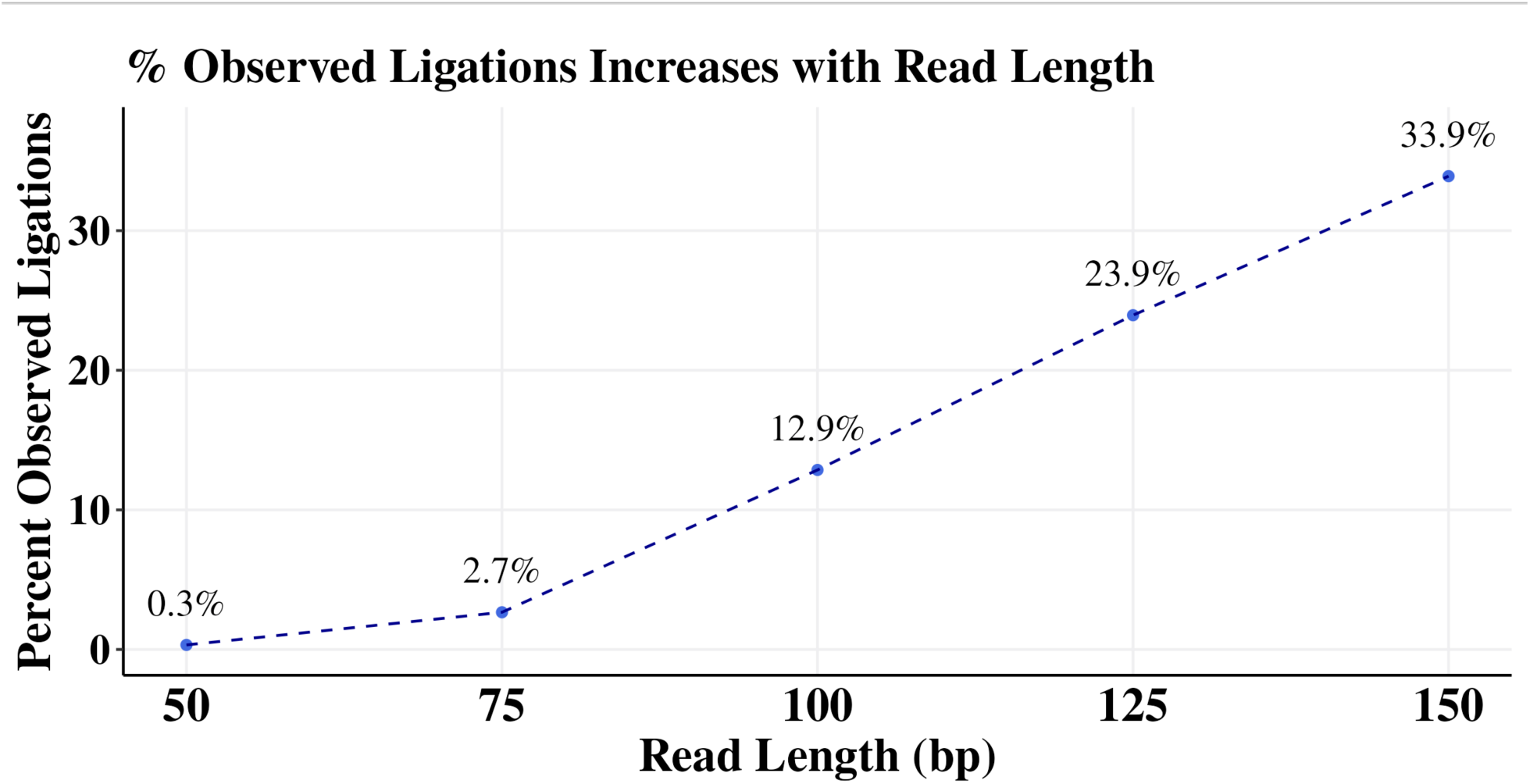
Percent observed ligations increases with read length.

**Supplementary Figure 2.**
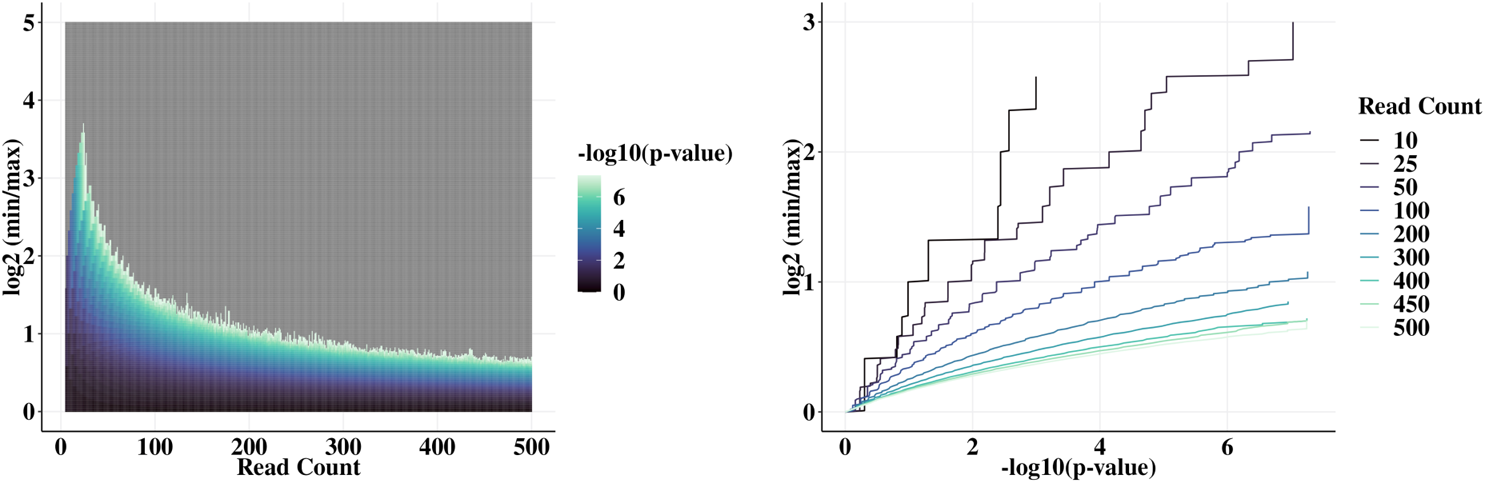
The probability of observing a high *FactorFinder* statistic under the null hypothesis is higher at low read counts.

**Supplementary Figure 3.**
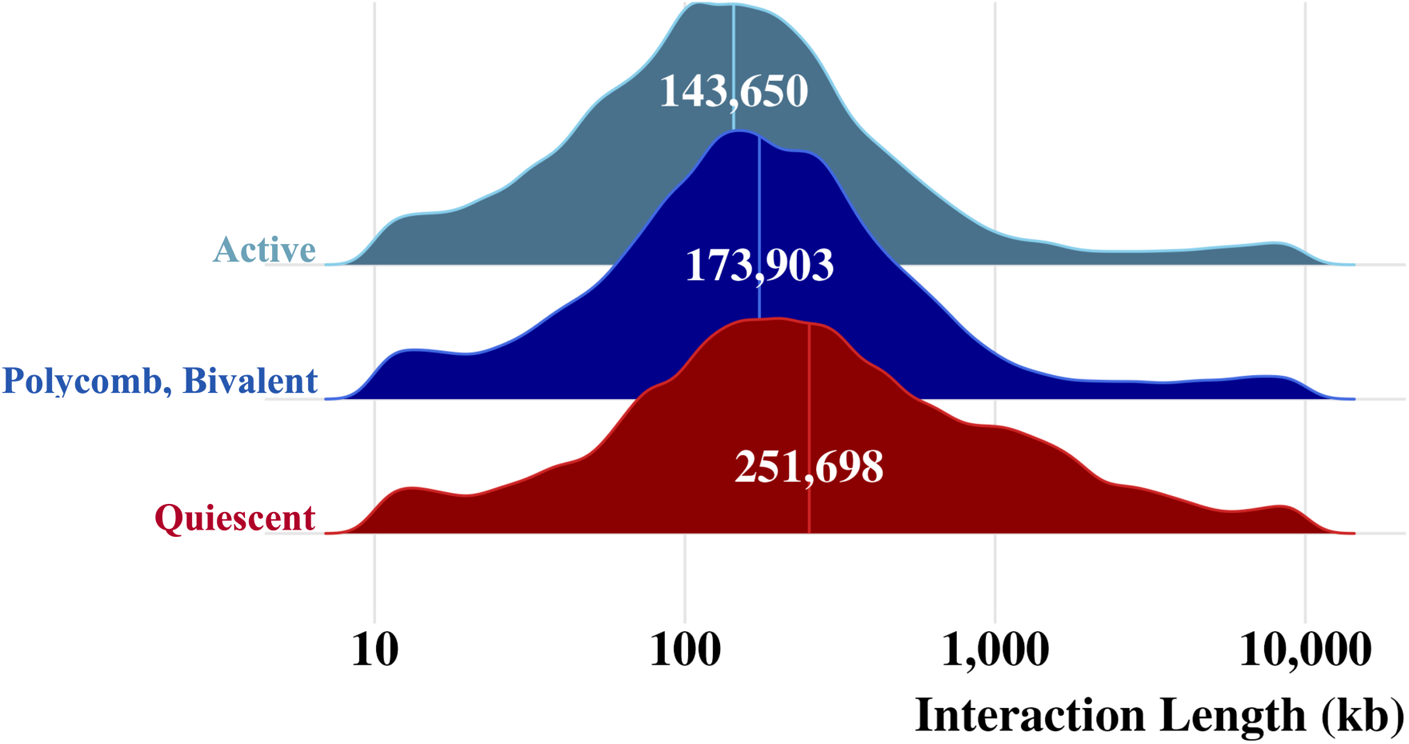
Cohesin extrudes significantly further through quiescent regions than active regions.

**Supplementary Figure 4.**
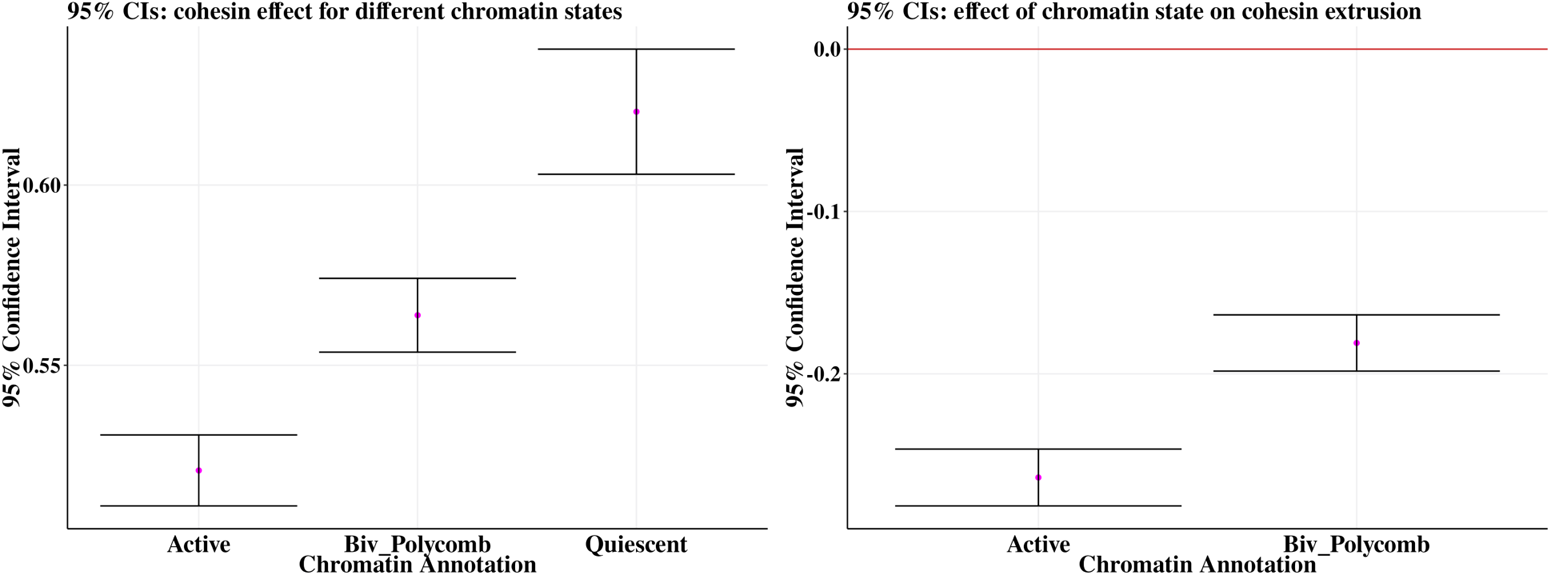
Controlling for locus-specific variation with linear mixed models does not attenuate the relationship between chromatin state and extruded loop size. Note that for the figure on the right, the group that active and bivalent polycomb are being compared to is quiescent.

**Supplementary Figure 5.**
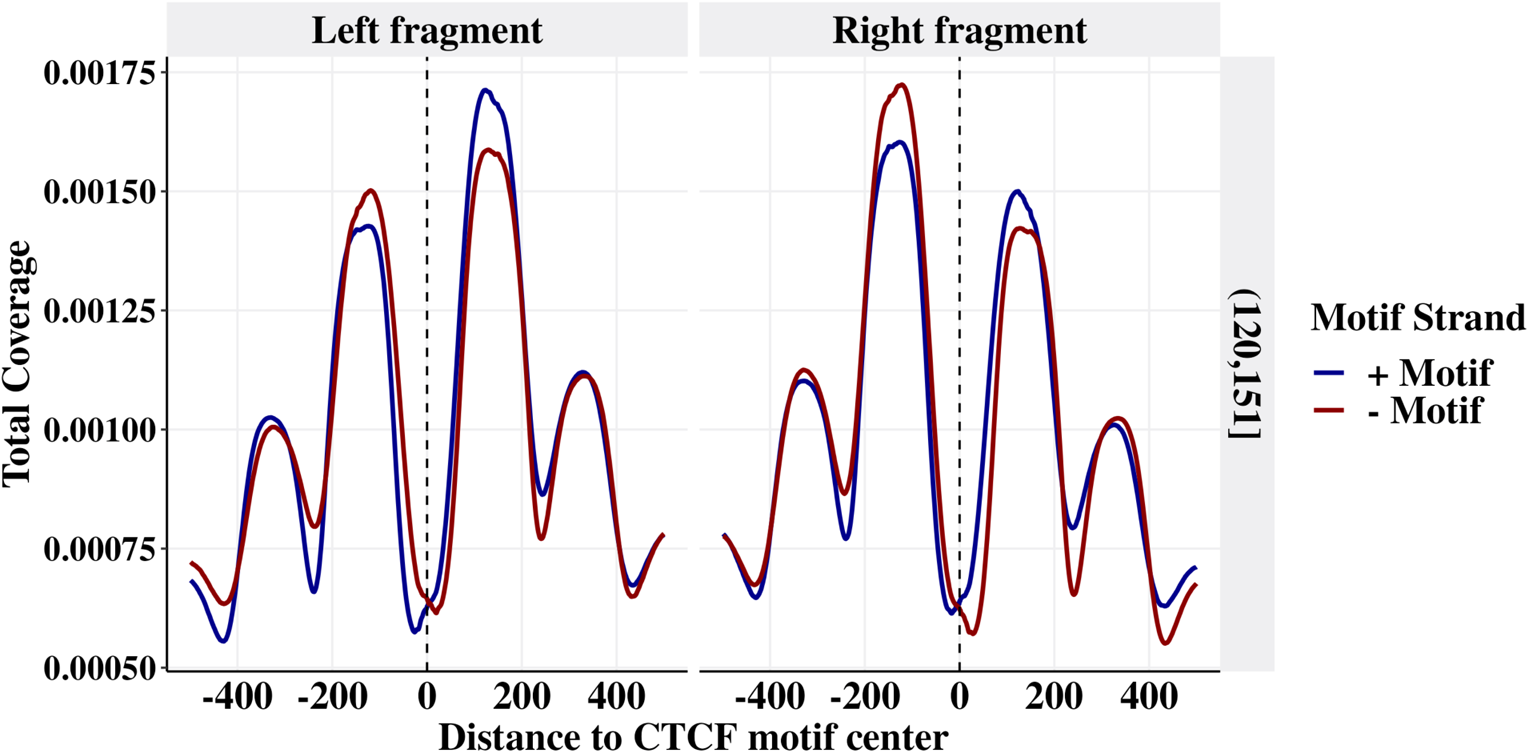
Nucleosomes are preferentially positioned inside the loop.

